# Unraveling the Half and Full Site Sequence Specificity of the *Saccharomyces cerevisiae* Pdr1p and Pdr3p Transcription Factors

**DOI:** 10.1101/2023.08.11.553033

**Authors:** Evan R. Buechel, Heather W. Pinkett

**Affiliations:** Department of Molecular Biosciences, Northwestern University, Evanston, IL 60208, USA

**Keywords:** Pdr1, Pdr3, Zinc cluster transcription factors, Pleiotropic drug resistance, CUT&RUN, DNA binding, Transcription

## Abstract

The transcription factors Pdr1p and Pdr3p regulate pleotropic drug resistance (PDR) in *Saccharomyces cerevisiae*, via the PDR responsive elements (PDREs) to modulate gene expression. However, the exact mechanisms underlying the differences in their regulons remain unclear. Employing genomic occupancy profiling (CUT&RUN), binding assays, and transcription studies, we characterized the differences in sequence specificity between transcription factors. Findings reveal distinct preferences for core PDRE sequences and the flanking sequences for both proteins. While flanking sequences moderately alter DNA binding affinity, they significantly impact Pdr1/3p transcriptional activity. Notably, both proteins demonstrated the ability to bind half sites, showing potential enhancement of transcription from adjacent PDREs. This insight sheds light on ways Pdr1/3 can differentially regulate PDR.

## INTRODUCTION

Zinc cluster transcription factors are a diverse family of proteins that are unique to fungi and play important roles in regulating gene expression in response to environmental cues(1). This family of transcription factors possesses three distinct domains: a DNA binding domain (DBD) composed of a Zn_2_Cys_6_ zinc finger (or binuclear cluster) and dimerization helices of coiled-coils, regulatory domain, and a highly charged activation domain. Zinc cluster transcription factors predominantly bind to DNA as dimers, recognizing two CGG triplets with varying spacing and orientation between repeats. The structural studies of several fungal zinc cluster proteins, including Gal4 in *Saccharomyces cerevisiae*, serve as a model for understanding structure and function of this family of transcription factors(2–6). These structures reveal how the Zn_2_Cys_6_ zinc finger recognizes a CGG nucleotide triplet via major groove contacts. Additional sequence specificity is imposed by interactions between the spacer nucleotide sequence and the flexible linker that connects the Zn_2_Cys_6_ zinc finger and dimerization helices(7).

Despite the wide range of spacing and orientations available to zinc cluster proteins, there are several *S. cereivisiae* zinc cluster proteins including Pdr1p, Pdr3p, Pdr8p, Rdr1p, and Yrr1p that recognize a zero-gap everted repeat (5’-CCGCGG-3’)(8–11). To date no structure has been solved of a zinc cluster protein with this binding orientation, which leads to uncertainty in understanding the protein-DNA interface. The absence of a nucleotide spacer between CGG triplets, suggests that additional sequence specificity may result from interactions with the nucleotides flanking the everted repeat. In addition, what is known about the current binding motifs for these zinc cluster proteins does not fully explain the variation in the genes they control, suggesting the existence of a sequence specificity mechanism that has yet to be fully characterized.

The most prominent members of the zero-gap everted repeat zinc cluster proteins are Pdr1p and Pdr3, the primary transcriptional regulators of the Pleiotropic Drug Resistance (PDR) network(12–17). The PDR network, involved in multidrug resistance in yeast, is also triggered by a variety of general stresses including mitochondrial damage, heat shock, membrane damage and translational stress(18–21). The expression of ATP-binding cassette (ABC) transporters and major facilitator transporters are strongly induced by PDR network activation and are responsible for export of a wide range of toxic compounds(22–27). Pdr1p and Pdr3p recognize specific PDR responsive elements or PDREs, with single or multiple PDREs located in the promoters of the PDR genes, including *SNQ2, RSB1, TPO1, PDR3* and the well-characterized promoter of ABC transporter *PDR5* (Figure 1A)(8,22,28–30). PDREs have been classified into four types, with type A being the canonical 5’-TCCGCGGA-3’(31). The other three types, B, C and D represent observed single base substitutions from the canonical sequence (Figure 1B). Although both proteins recognize PDREs in the promoters of PDR genes, not all PDRE-containing promoters are regulated by both proteins. This differential regulation of the PDR network by Pdr1p and Pdr3p is not yet fully understood, despite their significant role as the primary transcription regulators of this network.

**Figure 1.**
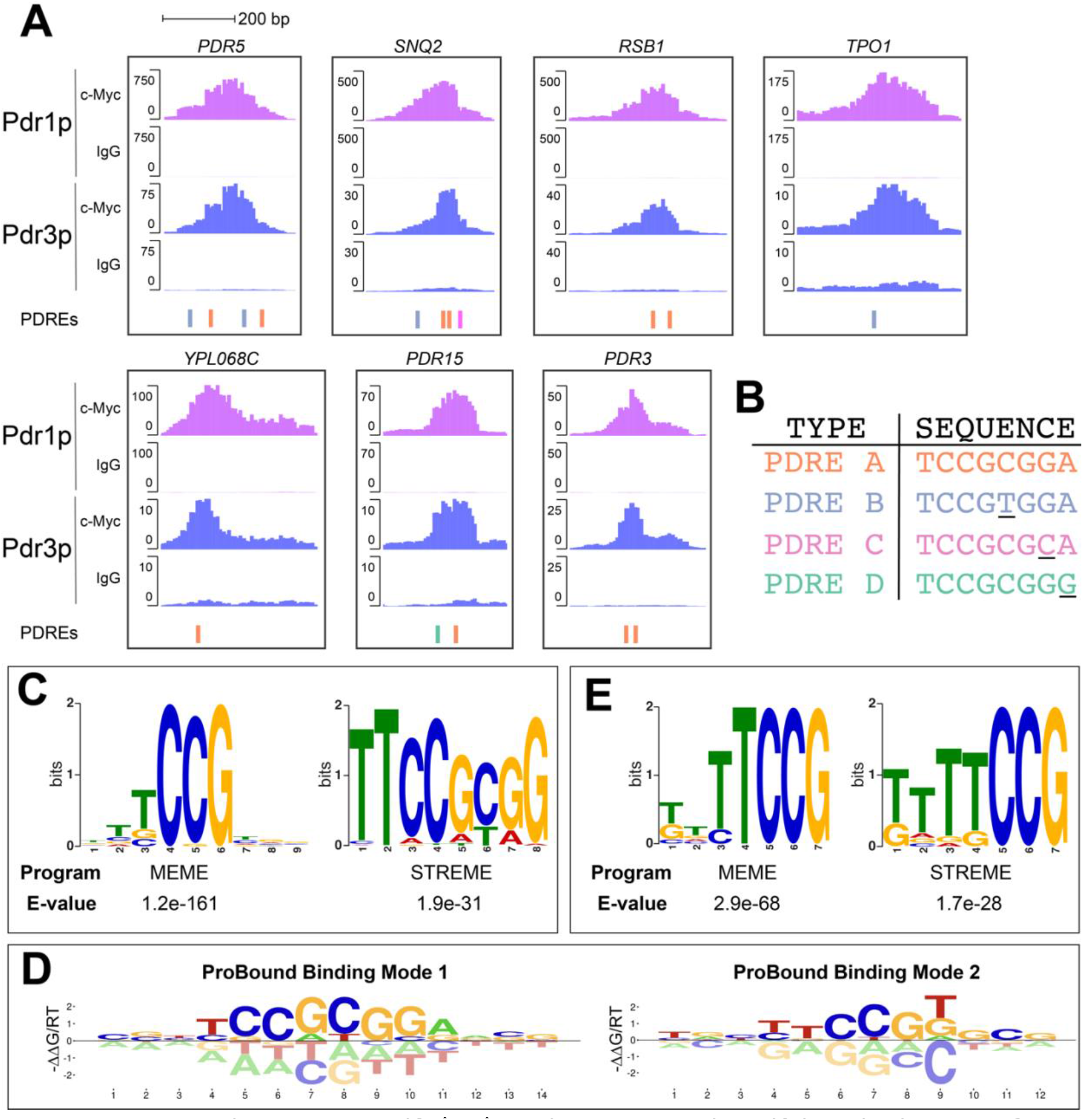
Pdr1p and Pdr3p bind PDRE motifs *in vivo*, with programs identifying binding to half and full PDRE sites. (**A**) Representative genome browser shots of the shared CUT&RUN Pdr1p and Pdr3p peaks at 7 gene promoters, with both samples normalized to their IgG negative controls. The location and type of PDRE is annotated for each peak. (**B**) Nucleotide sequence of the four PDRE types, where type B-D are one nucleotide substitution (underlined) from the canonical type A. (**C**) The top scoring de novo motifs discovered by MEME-ChIP from the Pdr1p CUT&RUN peaks. (**D**) First two binding modes fitted by ProBound trained on 500k Pdr1p CUT&RUN reads with symmetry constraints on binding mode 1. (**E**) Top scoring de novo motifs discovered by MEME-ChIP from Pdr1p CUT&RUN peaks that lack the four types of PDREs.

Previous experiments using ChIP-chip and ChIP-seq have shown few shared peaks for each protein, leading to uncertainty in the Pdr1p and Pdr3p regulons. To address this issue, a more sensitive, higher resolution technique for genomic occupancy was employed. Given that both Pdr1p and Pdr3p bind a gapless, everted CGG repeat, we propose that their in vivo binding specificity has additional dependency on flanking sequences beyond what has been previously described, leading to the observed differences in regulons. Here we use a combination of genome occupancy profiling, in vitro binding assays, and transcriptional reporter assays to investigate the role of DNA sequence on promoter occupancy, binding affinity and transcription. Our investigation uncovers differing specificity for sequence variation within PDREs for both Pdr1p and Pdr3p. Moreover, we find that flanking sequences play a significant role in PDRE function with some capable of altering the binding affinity by more than 2-fold and transcription by more than 10-fold. In addition, our findings provide evidence that these proteins recognize and have high affinity for half sites that contribute to the observed genomic occupancy. Collectively, this study provides unprecedented detail into the impact of binding site sequence variation on both DNA binding affinity and transcriptional activity of Pdr1p and Pdr3p.

## MATERIAL AND METHODS

### Construction of yeast strains

All strains were derived from the FY1679-28c parent strain. Strains FY1679-28c, ΔPDR1, ΔPDR3, ΔPDR1ΔPDR3, and YRP10 were a generous gift from Dr. Karl Kuchler. The strain YRP10 contains Myc-Pdr1p and HA-Pdr3p and has been described elsewhere along with ΔPDR1, ΔPDR3, and ΔPDR1ΔPDR3(32). To construct the Pdr3p-3xMyc strain, the tagging plasmid pTF277 (Addgene 44094) was altered by PCR to replace the 2xStrep tag with 3xMyc. The tagging cassette was PCR amplified with 40-nt flanking sequences for homologous recombination and transformed into FY1679-28c(33). Successful integration was verified by colony PCR, while protein function was confirmed by resistance to cycloheximide in a spot dilution assay(32). Yeast cells were grown at 30°C in YPD (1% w/v yeast extract, 2% w/v peptone, 2% w/v dextrose) unless otherwise noted.

### CUT&RUN

Yeast nuclei were purified as described previously by Orsi *et al*(34). In brief, yeast cells (YRP10 and Pdr3p-3xc-Myc) were grown at 30°C to an OD_600_ of 0.6-0.8, then spheroplasted with Zymolyase 100T (USBiological Z1004). The nuclei were isolated, flash frozen in liquid nitrogen and stored at -80°C. CUT&RUN assays were carried out using the Epicypher CUTANA ChIC/CUT&RUN Kit (Epicypher 14-1048). 1 million nuclei were thawed, immobilized on concanavalin A beads, and permeabilized in 0.05% digitonin cell permeabilization buffer. For each sample, the permeabilized nuclei were divided equally and incubated in 50 µL antibody buffer with either 0.5µg of control rabbit IgG antibody (Epicypher 14-1048) or anti-c-Myc antibody (ThermoFisher Scientific 13-2500) overnight at 4°C. After washing the beads, pAG-MNase (EpiCypher 14-1048) and 100mM CaCl_2_ was added to the immobilized cells and incubated for 2h at 4°C to digest chromatin DNA. DNA cleavage was terminated by addition of 150 µl of stop buffer and 0.5 ng of *E. coli* Spike-in DNA (EpiCypher 14-1048). After incubation for 10 minutes at 37°C, the CUT&RUN-enriched DNA was phenol-chloroform precipitated with 80 µg glycogen. The DNA was dissolved in 0.1X TE buffer, followed by DNA quantification using the Quant-iT PicoGreen assay (Thermofisher Scientific P7589). CUT&RUN libraries were prepared using the NEBNext Ultra II DNA Library Prep Kit for Illumina (NEB E7645S) following the manufacturer’s instruction with modifications to SPRI bead size selection and library amplification reactions to retain small DNA fragments(35). Amplified libraries were size selected with Sera-Mag Select (Cytiva 29343045) beads and quality assessed using the Agilent TapeStation HS DNA assay (Northwestern University NUseq core). CUT&RUN libraries were pooled and sequenced on the Illumina Nextseq 500 platform with 75bp paired end reads (Northwestern University NUseq core).

### Next generation sequencing analysis

Paired-end sequencing reads were trimmed with Trimmomatic to remove adaptor sequences from the 3’ end of each read(36). Then reads were aligned to the reference *S. cerevisiae* R64 assembly or the *E. coli* K12 assembly using Bowtie2 with settings --end-to-end,--dove-tail,--phred33(37). Picard was then used to sort, index and mark as duplicates the alignments using the MarkDuplicate function. The program deepTools was used to filter the resulting BAM files to remove sequences longer than 120bp(38). Peaks were called using MACS2 v2.1.1 with the settings –c –g ce –q 0.00001 --keep-dup all –f BAMPE –SPMR(39).

For visualization bigWig coverage files were generated by deepTools with samples normalized using the *E. coli* spike-in DNA. A scale factor based on the proportion of reads mapping to the *S. cerevisiae* or the spike-in genome was calculated(40). The bamCoverage function of deepTools was used to produced normalized bigWig files that were loaded into IGV-Web for peak visualization(41). Peak annotation was performed using the R package rGreat v1.99.5 in basalPlusExt mode(42). The resulting gene list was filtered to only include genes with a peak summit within -1000 to +500 of the transcription start site.

### De novo motif discovery

MEME-ChIP was used to perform a comprehensive motif analysis on CUT&RUN sequences(43). Sequences from –250 to +250 bp from the summits of all peaks called by MACS2 were analyzed for enrichment. MEME-ChIP was run with the setting -minw 5 -maxw 12 -bfile, where the background file is a 4^th^ order Markov model from yeast noncoding DNA(44).

Analysis of the CUT&RUN data by ProBound followed the default ChIP-seq protocol except for a modified preparation of the count table(45). To take advantage of the higher resolution CUT&RUN data, 120 bp centered on each alignment was used as input to build the ProBound count table rather than the default 200 bp downstream of the 5’ end of mapped reads. The count tables were submitted to the ProBound web server along with a JSON configuration file. Several configurations were tested, with the final results produced by 4 binding modes with the first mode seeded with ‘TCCGCGGA’ and symmetry constraints. Configurations were assessed by comparing the relative affinities predicted by the binding mode versus the experimentally measured affinities.

### DNA binding domains expression and purification

The DNA sequences encoding the DNA binding domains of Pdr1p (residues 1-207) and Pdr3p (residues 1-184) were amplified by PCR from the Yeast Genomic Tiling Collection (GE Healthcare YSC4613). The PCR products were inserted into the pMCSG7 vector by ligation independent cloning, then transformed into *E. coli* Rosetta pLysS(46). Cells were grown at 37°C to an OD_600_ of 0.3, then continued to grow at 16°C to an OD_600_ of 0.7. The cultures were induced with 0.5mM IPTG in the presence of 100µM Zinc acetate and continued to grow for 18 hours. The harvested cells were lysed in Buffer A (25mM Tris pH 7.5, 500mM NaCl, 10% glycerol, 5mM BME, 15mM imidazole) and 1mM PMSF (Phenylmethanesulfonyl fluoride) by sonication. The lysate was centrifuged at 17,000 rpm for 30 minutes and the supernatant applied to a Ni-NTA column. The column was washed with 10 column volumes (CV) of Buffer A, followed by 10 CVs Buffer A supplemented to 1M NaCl and 45mM imidazole. The bound protein was eluted with 5 CVs of Buffer A supplemented to 250mM imidazole. The pure protein was subjected to TEV cleavage overnight in Dialysis Buffer (25mM HEPES pH 7.5, 400mM NaCl, 10% glycerol) supplemented with 5mM BME. The cleaved protein was diluted to 100mM NaCl and clarified by centrifugation before application to a 5mL Heparin HiTrap HP (GE Healthcare 17-0407-01) column equilibrated with Buffer B (25mM HEPES pH 7.5, 100mM NaCl, 10% glycerol). The Heparin column was washed with Buffer B before a 0-50% gradient, followed by 100% Buffer C (25mM HEPES pH 7.5, 1000mM NaCl, 10% glycerol) was applied to elute the bound protein. Fractions that were judged by SDS-PAGE to be approximately 95% pure were pooled and dialyzed against Dialysis Buffer supplemented 0.5mM TCEP.

### Fluorescence polarization

All unlabeled and 5-FAM oligonucleotides were synthesized by Integrated DNA Technologies (IDT). Oligonucleotides were resuspended in ddH_2_O, then complementary strands were mixed at an equal molar ratio in 25mM HEPES pH 7.5 and 50mM NaCl. The mixed oligonucleotides were annealed by heating to 90°C for 5 minutes, before cooling to room temperature overnight. Annealed DNA was stored at -20°C. For binding assays, a 2-fold serial dilution of each protein was prepared in Dialysis Buffer. The binding buffer and incubation time at 4°C was optimized for both proteins. FAM-labeled DNA at a final concentration of 2nM was mixed with protein in a Greiner Bio-one 384-well plate (781900) in a final buffer of 25mM HEPES pH 7.5, 50mM NaCl, and either 10% glycerol for Pdr1p or 2.5% glycerol for Pdr3p. The plate was incubated at 4°C for 2 hours for Pdr1p, then equilibrated for 5 minutes at 18°C in a Spark (Tecan) plate reader prior to fluorescence polarization measurements. For Pdr3p the plate was immediately equilibrated at 18°C. For competitive binding experiments, 2-fold serial dilutions of unlabeled oligonucleotides were incubated with a constant concentration of protein and 2nM labeled DNA overnight at 4°C for Pdr1p, and 10 minutes at 4°C for Pdr3p. All binding data was measured using a Spark plate reader (λ_ex_ = 495nm, λ_em_ = 520nm). The data was analyzed and fit in Graphpad Prism 9.0 using a sigmoidal 4PL regression. Representative curves are shown for one experiment (three technical replicates) and were repeated at least two times in triplicate.

### Luciferase assay

All luciferase reporter plasmids were derived from pCA955, which was a generous gift from Dr. Claes Andréasson (47). Plasmid pCA955-NheI was constructed by PCR mutagenesis of pCA955 to replace the heat shock element in the minimal CYC1 promoter with a NheI restriction site. IDT synthesized oligonucleotides for each PDRE were phosphorylated, annealed, and then ligated into NheI linearized pCA955-NheI. Ligated plasmids were transformed into *E. coli* Top10 competent cells. Successful transformants were verified by sequencing, then transformed into yeast strains by the Lithium Acetate method(33).

Yeast cultures were grown overnight at 28°C in synthetic growth medium containing 0.67% yeast nitrogen base and 2% dextrose, supplemented to support the growth of auxotrophic strains. The following morning, while in exponential phase, cultures were adjusted to the same cell density of 0.5 OD and transferred to white 96-well plates (Corning 3917). Nano-Glo substrate (Promega N1110) was diluted 1:100 with the supplied lysis buffer and mixed 1:10 with the cells in the white 96-well plates. Bioluminescence was measured in a Spark (Tecan) plate reader every two minutes for an hour at 28°C. Each culture growth and luciferase assay were repeated two additional times on separate days. Since bioluminescence plateaus at 20 minutes, all data was analyzed at this timepoint. The luminescence of the empty reporter plasmid (pCA955-NheI) in each strain was subtracted from all reporter plasmids in the same strain. All data was analyzed in Microsoft Excel and GraphPad Prism 9.0.

## RESULTS

### Genomic occupancy of Pdr1p and Pdr3p in *S. cerevisiae* using CUT&RUN

We profiled the genomic occupancy of Pdr1p and Pdr3p in *S. cerevisiae* using CUT&RUN, which allows high resolution detection of transcription factor binding sites(40). Based on the CUT&RUN data, it appears that both Pdr1p and Pdr3p bind most strongly at the promoters of seven genes, all of whom are known to be strongly regulated by these transcription factors, including *PDR5*, *SNQ2*, *TPO1*, *RSB1*, and *PDR3* (Figure 1A)(31). However, while Pdr3p was limited to seven genes, Pdr1p has substantial additional peaks. In addition to the shared PDR genes, there is another 86 Pdr1p peaks that contain any of the four PDRE types (Figure 1B). These peaks are closely centered on the PDREs, with the median peak summit within 29bp of the PDREs. As anticipated, the number of PDREs in the Pdr1p peaks is correlated (Pearson r = 0.65, r^2^ = 0.42) with the peak signal (Sup. Figure 1A). However, 42 of Pdr1p peaks without PDREs possess a peak signal greater than the median of the 1 PDRE peaks, indicating that Pdr1p tolerates more divergent sequences than the types A-D PDREs. Within the single PDRE peaks the type C PDREs were the least abundant and had significantly lower peak signal than the type A, B, and D PDREs (Sup. Figure 1B). Unlike the previous ChIP-Seq experiment by Fardeau and colleagues, we observe peaks containing just a type C or D PDRE indicating that type A and B PDREs are not required for binding(48). Across all Pdr1p peaks, type B PDREs were the most common, followed by type A and D, while type C were less frequent (Sup. Table 1). We also observe that a number of previously identified PDREs were either not bound or very weakly bound by Pdr1p in the CUT&RUN dataset. Some of these unbound PDREs are located in promoters that have previously been described as directly regulated by Pdr3p but not Pdr1p, such as the promoter region of one of the PDR major facilitator transporters *FLR1*(26,49,50).

**Table 1.**
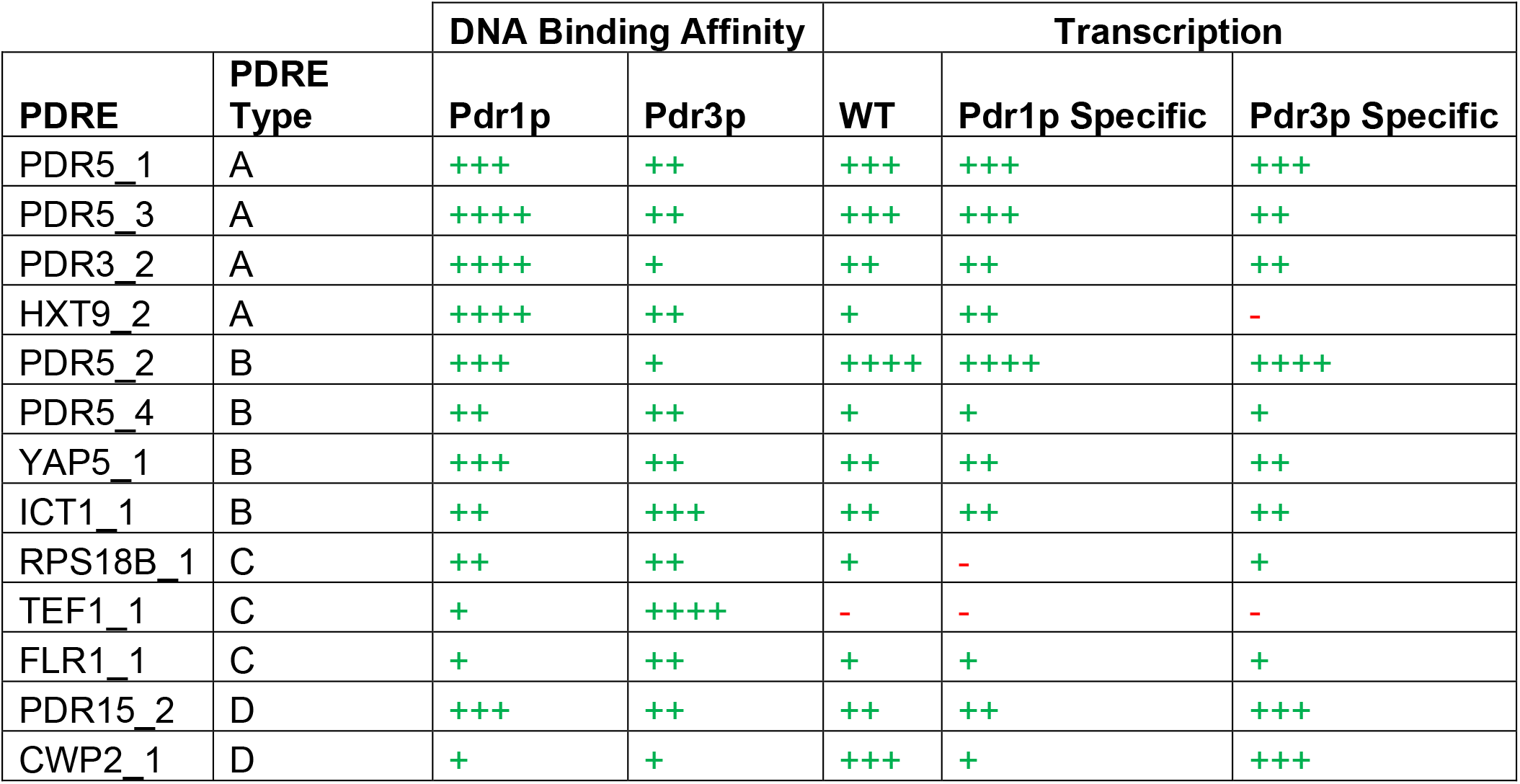
Summary of DNA binding affinity and transcriptional activity of all tested PDREs. For each category PDREs were grouped into quartiles (-, +, ++, +++, and ++++ correspond to 0%, 0-25%, 25-50%, 50-75%, and 75-100% of maximum value or tightest affinity). Transcriptional activity was calculated for total (WT strain), Pdr1p specific (ΔPDR3 strain - ΔPDR1ΔPDR3 strain), and Pdr3 specific (WT strain - ΔPDR3 strain).

The vastly fewer Pdr3p CUT&RUN peaks in our datasets (7 versus 792) is likely due to the approximately 10-fold lower abundance of Pdr3p to Pdr1p in logarithmic growth(51). This aligns with previous studies that highlighted that Pdr3p expression is regulated by Pdr1p and Pdr3p through the presence of two PDREs in the *PDR3* promoter, in agreement with the strong binding to the *PDR3* promoter by both transcription factors in CUT&RUN(Figure 1A)(52). Conditions that increase Pdr3p expression, like loss of normal mitochondrial function, would likely reveal occupancy of additional sites that align with the known Pdr3p regulon(18).

### Identification of Pdr1p PDRE flanking specificity from CUT&RUN data

Despite the presence of known PDRE sites, the occupancy level at sites with identical core sequences in our CUT& RUN data differs drastically, suggesting that core sequence alone is not sufficient to dictate strong binding. Lack of genomic occupancy at previously identified PDREs led us to explore the possibility of refining the published PDRE motifs to incorporate any tolerated variation in the core PDRE and any additional flanking sequence specificity that would better classify the genomic binding sites. To achieve this, we used MEME-ChIP program suite and the recently published ProBound to do de novo motif discovery(43,45).

The 792 Pdr1p peaks were submitted to MEME-ChIP using both MEME and STREME for motif discovery, followed by CentriMo to evaluate central enrichment. The top-scoring MEME motif was the core CGG triplet, which has been observed for other zinc cluster transcription factors. The top scoring STREME motif was the PDRE-like 5’-TTCCGCGG-3’ (Figure 1C). Both motifs were found to be strongly enriched within the center 140bp of the Pdr1p peaks, indicating that both motifs are associated with Pdr1p rather than cofactors.

The inability to produce a PDRE motif with MEME-ChIP that both captures the different PDRE types and more flanking sequence led us to explore whether other motif discovery tools could capture additional sequence specificity. The recently published ProBound algorithm uses machine learning to produce binding modes that capture protein binding preferences from Next Generation Sequencing (NGS) data. Unlike MEME-ChIP, ProBound’s binding models represent the energetic contribution of each base to binding rather than a position weight matrix. ProBound was trained with 500k reads randomly sampled from the Pdr1p CUT&RUN data, and a primary symmetric binding mode was seeded with the canonical PDRE 5’-TCCGCGGA-3’. In addition, three additional binding modes were included to capture any alternate motifs such as cofactors (Figure 1D). The resulting primary binding mode generated by ProBound spans 14bp, including two additional nucleotides on each flank, making it wider than the 10bp MEME PDRE motif. Moreover, the PDRE ProBound model takes into account the tolerated mutations in the PDRE that correspond to types B-D PDREs.

The second Probound binding model, which is centered on a single CCG triplet, was found to correlate with the top scoring motif produced by MEME-ChIP (Figure 1C-D). Based on the similarity of these two motifs, we decided to investigate whether these motifs could represent half site binding of Pdr1p. We used the MEME suite tool FIMO to identify the Pdr1p peaks containing a PDRE, then repeated motif discovery with MEME-ChIP on the PDRE-free peaks. Removal of the PDRE containing peaks produced a more refined motif centered on a CCG triplet, suggesting that this represents a secondary binding motif that is distinct from the PDRE full site binding (Figure 1E). A search in the literature for a similar binding motif revealed a Pdr1p protein binding microarray (PBM) experiment by Badis et al. and a PBM experiment for Pdr3p by Gordân et al. that both produced several top-scoring 8-mers containing a single CCG triplet(53,54). The top scoring non-PDRE 8mer, TTCCGGAA, is in close agreement with the second ProBound model and the CCG motif produced by MEME, suggesting that Pdr1p can bind to DNA at these half sites in vivo. The use of multiple motif discovery tools and correlation with motifs previously reported provides support for their potential biological significance.

### PDRE flanking sequences alter Pdr1p and Pdr3p binding affinity and activity

To gain more information on the sequence specificity of Pdr1p and Pdr3p, we expressed and purified the DNA binding domains (DBDs) of both proteins (Sup. Figure 2). We first looked to examine the binding of both proteins to the four PDREs in the well-characterized *PDR5* promoter that depends equally on both Pdr1p and Pdr3p for expression(22,28). The PDR5 promoter contains two types of PDREs, A sites with the canonical PDRE sequences 5’-TCCGCGGA-3’ (present in PDR5_1 and PDR5_3) and the 1 base substitution PDRE B sites 5’-TCCG<u>T</u>GGA-3’ (present in PDR5_2 and PDR5_4) that are the most commonly described classes of PDREs(31). We incubated both DBDs with fluorescently labeled DNA 18mers containing the 8bp PDREs and 5bp flanking sequences and measured their binding by changes in fluorescence polarization (Figure 2A-C). The results indicate significant differences in binding between Pdr1p and Pdr3p for the same 18-mer sequence across all four PDREs. Pdr1p exhibited considerably tighter binding (52-118nM) compared to Pdr3p (173-284nM). Among the four *PDR5* PDREs, Pdr1p’s binding affinity showed the following trend from tightest binding to weakest: PDR5_3>PDR5_2=PDR5_1>PDR5_4. Within the type A PDREs, PDR5_3 displayed a 1.76-fold tighter binding compared to PDR5_1, while within the type B PDREs, PDR5_2 had a 1.33-fold tighter binding affinity than PDR5_4, despite both pairs sharing a conserved core sequence. Notably, PDR5_4 exhibited a significantly steeper binding curve for Pdr1p compared to the other three PDREs.

**Figure 2.**
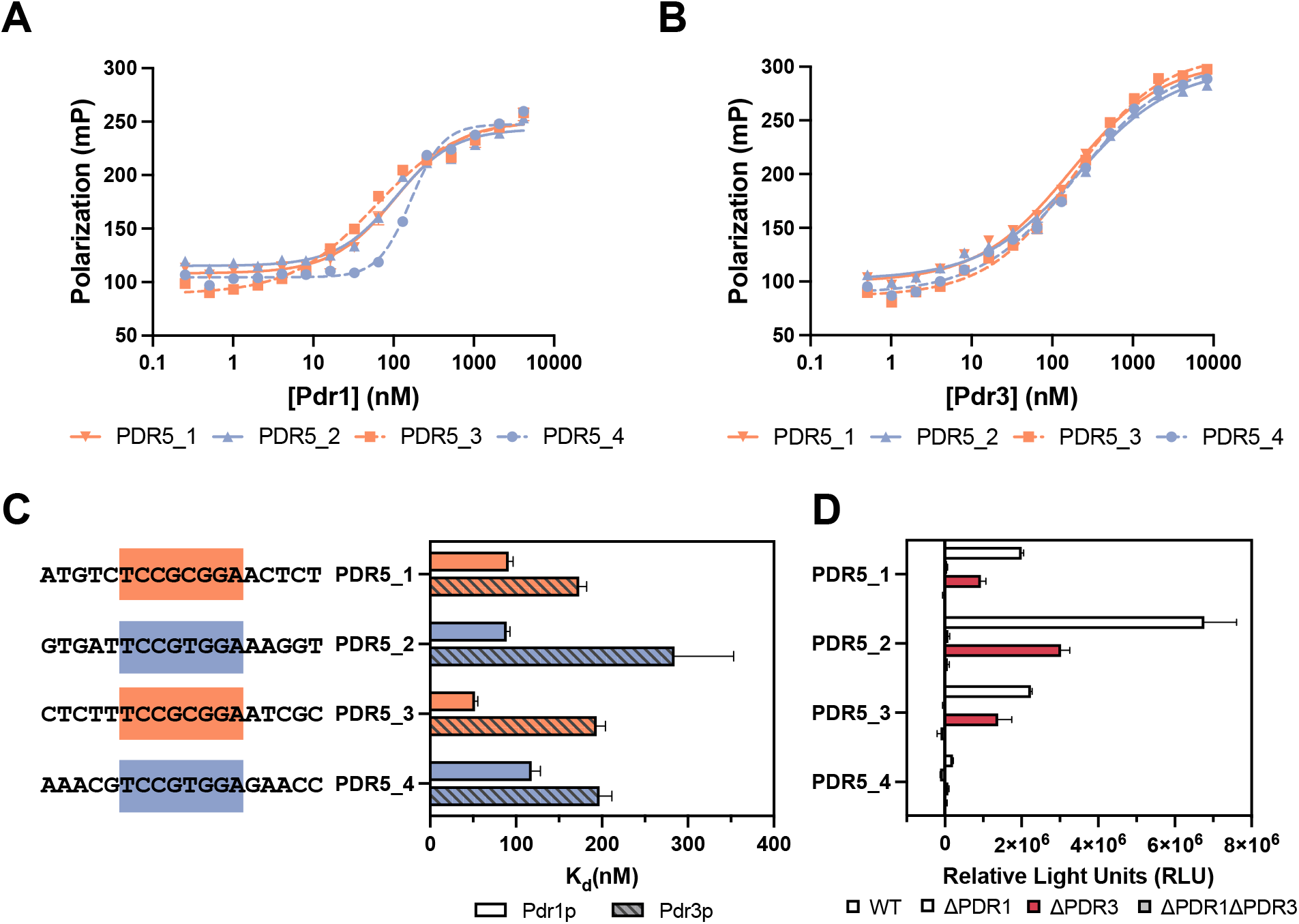
The *PDR5* PDREs are recognized with varying affinity by Pdr1p and Pdr3p despite their similar role in *PDR5* regulation. Fluorescence polarization binding curves of FAM-labeled *PDR5* PDRE DNAs incubated with 2-fold serial dilutions of either the (**A**) Pdr1p or (**B**) Pdr3p DBD. The 18mer DNAs encompass the *PDR5* PDREs with their flanking sequences. The data was analyzed in Graphpad Prism 9.0 using a sigmoidal 4PL fit. (**C**) Dissociation constants for the binding of Pdr1p and Pdr3p to the *PDR5* PDREs from the fitted fluorescence polarization binding curves, with the core PDRE sequence highlight by type. Representative curves are shown for one experiment (three technical replicates). (**D**) Transcription induction of luciferase reporter plasmids containing one of the 18mer *PDR5* PDREs inserted into a minimal *CYC1* promoter. Data points in panels C and D show standard deviations as error bars (n =3).

For Pdr3p, the hierarchy of binding strength from strongest to weakest follows the order of PDR5_1>PDR5_3=PDR5_4>PDR5_2. Interestingly, Pdr3p displayed the opposite preference for PDREs within each type, with PDR5_1 binding 1.12-fold tighter than PDR5_3 and PDR5_4 binding 1.44-fold tighter than PDR5_2. Both Pdr1p and Pdr3p demonstrated the strongest binding affinity to a type A PDRE and the weakest binding to a type B PDRE. However, their specificities for PDREs within each PDRE type differed. Notably, earlier analysis of promoter deletions in *PDR5* indicated that while PDR5_4 has no impact on transcription, the other three PDREs contribute equally to transcription(8,30).

In order to examine the relationship between binding affinity and transcription of individual PDREs, we introduced the 18mer PDRE sequences into a minimal *CYC1* promoter in a luciferase reporter plasmid. The transcriptional activity of all four *PDR5* PDREs was assessed in both a wild-type yeast strain and a strain with *PDR1* and/or *PDR3* deletions (Figure 2D). Consistent with previous analysis of the *PDR5* promoter, PDR5_4 exhibited significantly lower transcriptional activity compared to the other three PDREs. This suggests that the reduced activity of PDR5_4 is not due to increased distance from the transcriptional start site or other promoter context, but rather it is inherent to the PDRE itself with PDR5_4 being an intrinsically weak PDRE. PDR5_1 and PDR5_3 displayed similar levels of transcriptional activity, while PDR5_2 is 2-fold greater. Deletion of *PDR1* resulted in a near-complete loss of transcription from all four PDREs. This outcome can be attributed, at least partially, to the regulation of *PDR3* by Pdr1p(52). Interestingly, when *PDR3* is deleted, approximately half of the transcriptional activity for all four PDREs is lost. This aligns with the notion that the basal expression of PDR5 relies equally on both Pdr1p and Pdr3p(28).

Despite having the same core PDRE sequence, distinct binding patterns of Pdr1p and Pdr3p to PDR5_4 and PDR5_2 and the vastly different transcriptional activity indicates the flanking sequences play an essential role in binding site function. To investigate this hypothesis, we conducted experiments measuring DBD binding to various 18mer DNAs of PDR5_4. These sequences contained one or more base substitutions in the flanking region to more resemble the higher affinity PDR5_2 sequence (Figure 3, Sup. Figure 3). The results demonstrate that altering the flanking sequences of the PDR5_4 to resemble those of PDR5_2 bases leads to an increase in binding affinity for Pdr1p. This impact on binding affinity is particularly pronounced when single substitutions are introduced in base pairs immediately adjacent to the PDREs. However, even substitutions 3bp away from the core PDRE still affect affinity (Sup. Figure 3A). The majority of substitutions result in increased affinity, except for the unfavorable substitution of the 2^nd^ flanking 5’ base C4A. Specifically, the binding affinity is enhanced when either of the first two 5’ flanking bases are changed to T or the two 3’ flanking bases were changed to A. For instance, in the sequence 5’-AAAC<u>T</u>TCCGTGGA<u>A</u>AACC-3’ substituting the 1st flanking 5’ base G4T and the 3’ base G14A significantly increases affinity, although it is fully not sufficient to recapitulate the affinity of PDR5_2 (Figure 3A, 3C). However, when the unfavorable single substitution of C4A in the 2^nd^ flanking 5’ base is included with G4T and G14A substitutions, the affinity further increases, resulting in a sequence with an affinity that closely mirrors PDR5_2 (Figure 3A). This nonlinearity of base substitutions suggests that there is some level of context dependency that impacts Pdr1p binding, such as DNA shape (55,56). The difference in binding between Pdr3p and PDR5_4 versus PDR5_2 was minimal, making it difficult to assess the individual impact of substitutions (Sup. Figure 3D), However, similar to Pdr1p, the triple substitution of C4A, G4T, and G14A was enough to achieve the same level of affinity as PDR5_2 (Figure 3B). For both proteins the affinity for a sequence appears to be primarily determined by a 12bp sequence that includes the 8bp core PDRE and 2bp flanking nucleotides.

**Figure 3.**
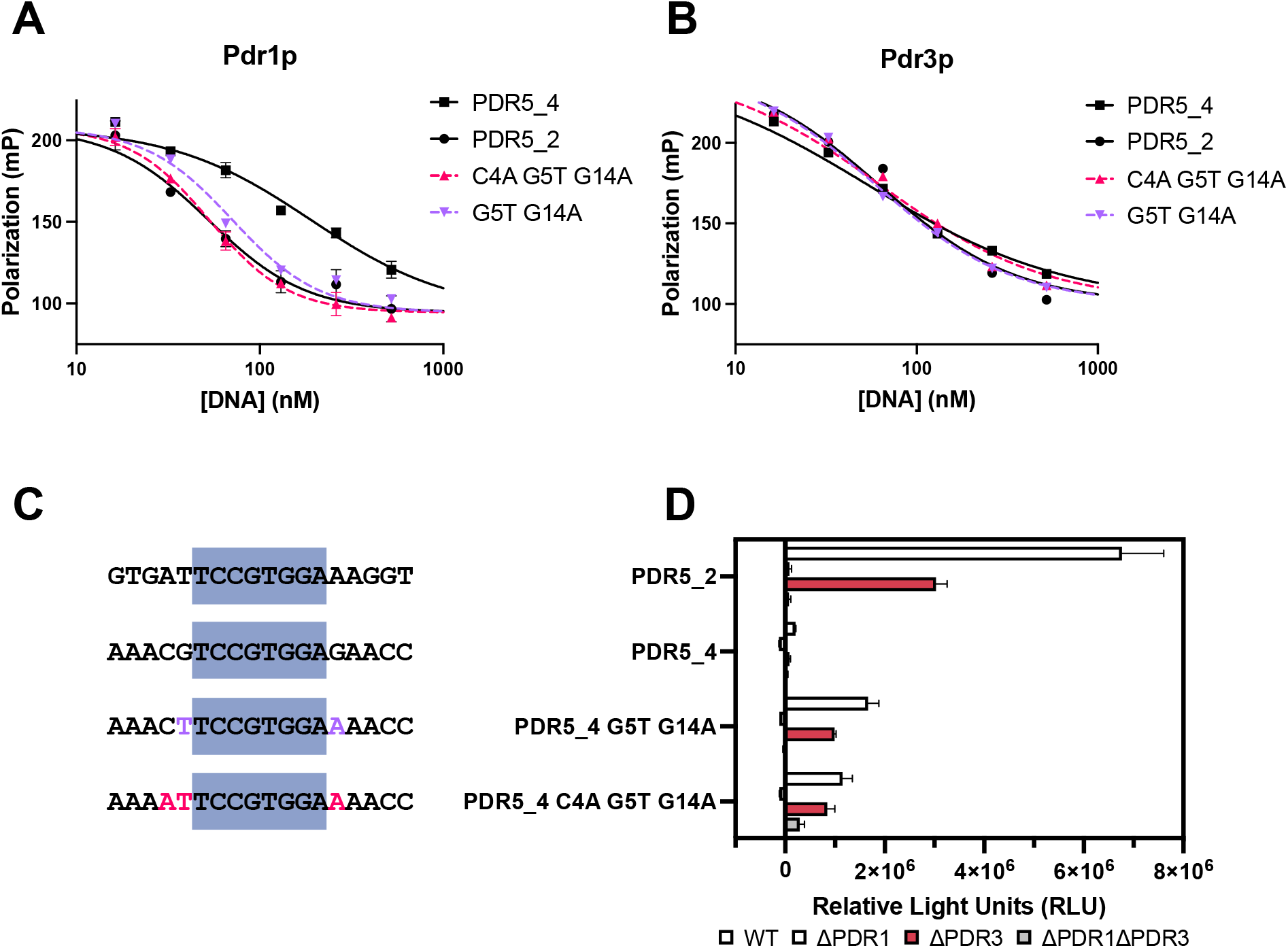
The modification of the *PDR5* PDRE flanking sequences alone alters both DNA binding affinities and transcriptional activity of Pdr1p and Pdr3p. The type B PDREs PDR5_2 and PDR5_4 were selected due to differences in both binding affinities and variable flanking sequences. Competitive fluorescence polarization binding curves of FAM-labeled PDR5_4 PDRE DNA and Pdr1p DBD (**A**) or Pdr3p DBD (**B**) incubated in the presence of unlabeled PDR5_4 PDRE DNA with either double or triple flanking base substitutions. Bases in the flanking sequences of the PDR5_4 PDRE were substituted with the corresponding base from PDR5_2 (**C**). The data was analyzed in Graphpad Prism 9.0 using a sigmoidal 4PL fit with a shared top and bottom for all datasets. Representative curves are shown for one experiment (three technical replicates) and were repeated at least two times in triplicate. **D**) Comparison of the transcription induction of luciferase reporter plasmids containing one of the WT 18mer *PDR5* PDREs or a PDR5_4 with substitutions inserted into a minimal *CYC1* promoter. Data points in panel D show standard deviations as error bars (n =3).

The favored substitutions to the PDR5_4 PDRE correlate with the flanking sequences in the motifs produced by MEME-ChIP and ProBound demonstrating that Pdr1 in vivo binding has a similar flanking sequence preference. The preferred 1st flanking nucleotides for Pdr1p is A, matching the extended consensus PDRE 5’-<u>T</u>TCCGCGGA<u>A</u>-3’(8). Among the models employed, the ProBound model in particular was able to capture the observed flanking sequence preferences for Pdr1p, with it correctly predicting the relative change of all the PDR5_4 single substitutions except for G14A. While the impact was lower, substitutions at the 3^rd^ flanking bases either 5’ or 3’ were able to alter affinity. This suggests that the 14bp wide PDRE binding mode produced by ProBound provides a reliable approximation for the full sequence that contributes to Pdr1p PDRE binding. Given that only a few substitutions were required for PDR5_4 to match the affinity of PDR5_2, we were curious about the effect of these substitutions on transcriptional activity. The double and triple substitutions in PDR5_4 that lead to high affinity, also significantly increase transcriptional activity (Figure 3D). However, the transcriptional activity of PDR5_2 remains considerably higher than either of the PDR5_4 substitution. This indicates that that affinity alone is not sufficient to achieve high levels of activity, emphasizing the substantial influence of the more distal flanking sequences on in vivo function of PDREs.

### Pdr1p and Pdr3p have different specificities for PDREs

We next wanted to assess whether the impact of flanking sequences on Pdr1p and Pdr3p binding affinity and activity is applicable to all PDREs or specific to the PDRE B sites. We selected additional sequences using three criteria: each type of PDRE must be represented, the flanking sequences must vary, and the sites were highly occupied in the Pdr1p CUT&RUN data. We also included the PDRE from the *FLR1* promoter as an example of a Pdr3p dependent site. The transporter *FLR1* contains a single type C PDRE in its promoter and has previously been shown to be directly regulated by Pdr3p but not Pdr1p(26,49,50). Investigation of the binding of Pdr1p and Pdr3p DBDs to these PDREs revealed that the two transcription factors do not share the same preferences for PDRE types. The results indicate that Pdr1p has a higher affinity for canonical type A PDRE, followed by type B, while C and D have weaker affinities and are more sensitive to changes in flanking sequences (Figure 4A, 4C). In contrast, Pdr3p has a stronger affinity for type C PDREs, followed by type B with the weakest binding to the type D and A PDREs (Figure 4B-C). For type B and C PDREs there is a marked difference in flanking sequence preference between Pdr1p and Pdr3p.

**Figure 4.**
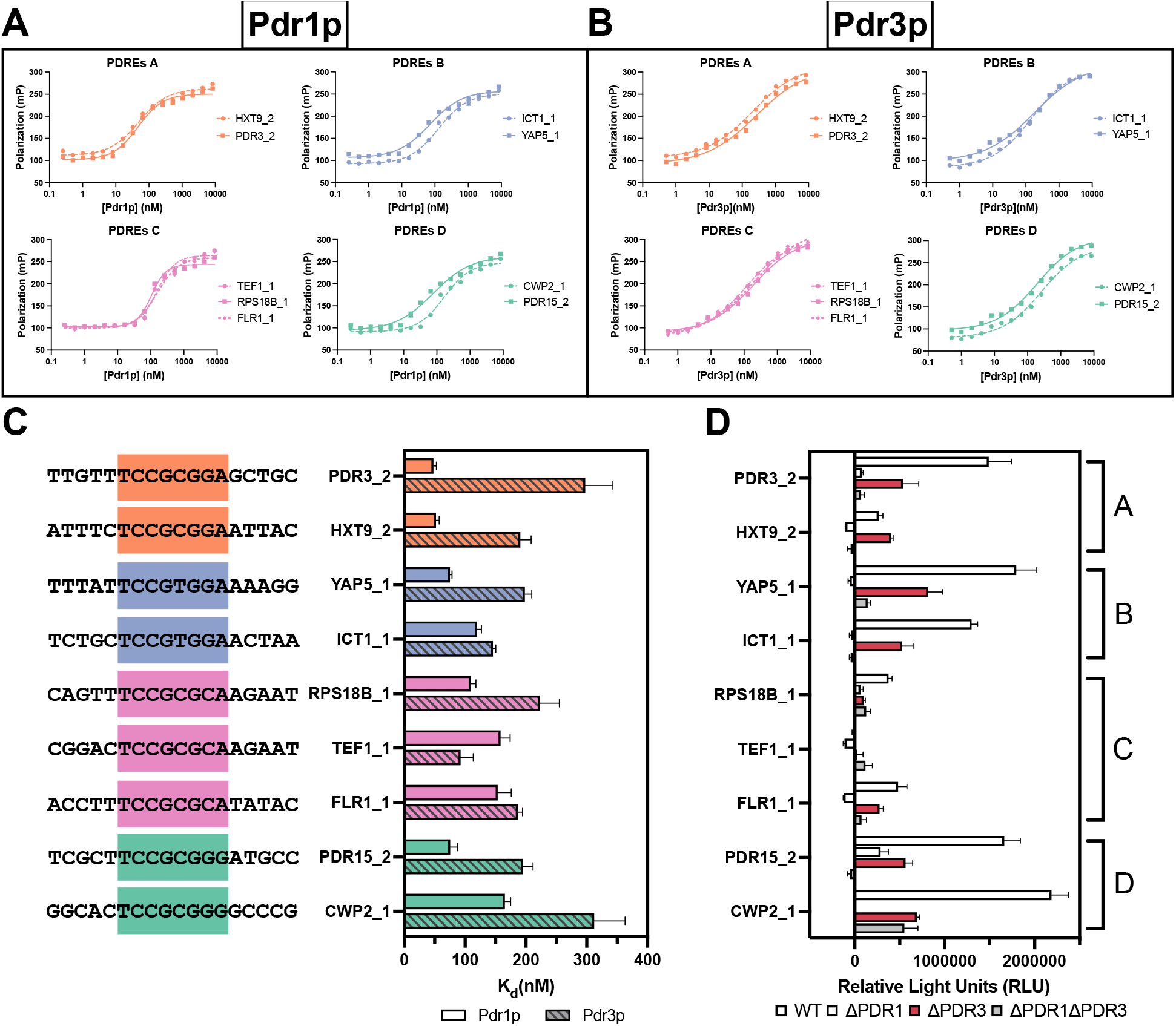
Pdr1p and Pdr3p have differing binding preferences for both PDRE types and flanking sequences. Variation within both the core PDRE or flanking sequences modulate transcriptional activity. Two of each PDRE type with varying sequences were selected from promoters bound in the Pdr1p CUT&RUN data, including some that are anticipated to be bound by Pdr3p. (**A, B**) Fluorescence polarization binding curves of FAM-labeled PDREs DNAs incubated with 2-fold serial dilutions of either the Pdr1p or Pdr3p DBD. The data was analyzed in Graphpad Prism 9.0 using a sigmoidal 4PL fit. (**C**) Dissociation constants for the binding of Pdr1p and Pdr3p to the PDRE DNAs from the fitted fluorescence polarization binding curves, with the core PDRE sequence highlighted by type. Representative curves are shown for one experiment (three technical replicates). (**D**) Transcription induction of luciferase reporter plasmids containing one of the 18mer PDREs inserted into a minimal *CYC1* promoter. Data points in panels C and D show standard deviations as error bars (n =3).

To assess the transcriptional activity of the different PDREs, a luciferase reporter assay was conducted (Figure 4D). Among the PDRE types, Type C PDREs exhibited weaker affinity compared to the other three types, particularly TEF1_1, which showed virtually no transcriptional activity. Interestingly, despite indications that *FLR1* is not directly regulated by Pdr1p, the FLR1_1 PDRE demonstrates Pdr1p-specific activity outside of its native promoter context. Similar to the observations with PDR5_2 and PDR5_4, the types A and B PDREs displayed a broad range of transcriptional activity. For the types A, B, and C PDREs, both Pdr1p and Pdr3p contribute roughly equally to transcriptional activity, except for HXT9_2 which is Pdr1p dependent (Table 1). On the other hand, the two type D PDREs appear to be more Pdr3p dependent. Notably, Pdr1p has minimal direct activity on the CWP2_1 PDRE, based on the minor difference in transcription between the ΔPDR3 and ΔPDR1ΔPDR3 strains. In the case of the PDR15_2 PDRE, it was the only PDRE tested that exhibited significant transcription in the ΔPDR1 strain and heavily reduced transcription in the ΔPDR3 strain. This finding aligns with *PDR15* mRNA levels primarily relying on the presence of Pdr3p(57).

The relationship between DNA binding affinity, genomic occupancy, and transcriptional activity for each PDRE was evaluated for Pdr1p (Sup. Figure 4). The ProBound binding mode generated from genomic occupancy data was able to predict high versus low affinity for PDRE sequences except for PDR5_1 (Sup. Figure 4A). Both the experimental binding affinities of Pdr1p and the ProBound predicted binding affinities showed a moderate correlation (R^2^ =0.4 vs 0.54) with the observed transcriptional activity of Pdr1p in the luciferase assay (when PDR5_2 is excluded) (Sup. Figure 4B-C). The analysis indicates the strength of DNA binding is not sufficient to accurately predict transcriptional activity, indicating the importance of other factors. Since the transcriptional activity of PDREs were measured in the absence of the native promoters, common factors such as nucleosome occupancy or presence of cofactors are unlikely involved in the observed differences. One possibility is that variation in the core and flanking PDRE sequence impacts not only DNA binding affinity but also protein conformation which in turn influences the level of transcription when bound.

### Pdr1p and Pdr3p recognize half site sequences

In addition to the CCGCGG triplet binding motif, a single CGG triplet binding motif was identified by both ProBound and MEME-ChIP from PDRE-free Pdr1p CUT&RUN peaks. This motif had been reported previously from PBM experiments, but no affinity dissociation constants or function was reported to date(53,54). To characterize the binding of Pdr1p and Pdr3p to a half site sequence for comparison to full site PDREs, we selected one of the top scoring 8mers (TTCCGGAA) from these two PBM experiments to test in vitro. To produce an 18mer sequence all the 8mers from the PBM experiments containing a single CGG triplet were examined to select favorable 5’ and 3’ flanking sequences, confirming that the remaining bases selected do not contain additional CGG triplets. This half site sequence with 5 nucleotide wide flanks was bound by Pdr1p at 229nM (Figure 5A-B). The affinity is weaker than the tested PDREs, but only slightly weaker than the type D PDREs. Mutating the core sequence to 5’-TTC<u>AA</u>GAA-3’ decreases affinity by 30-fold, which abolishes the CGG triplet, indicating binding specificity for this site is heavily dependent on the CGG triplet (Sup. Figure 5B). We tested a number of single base substitutions based on our discovered motifs but found little impact except for 5’-TTCCG<u>A</u>AA-3’ which increased affinity (Sup. Figure 5A). Pdr3p was also capable of binding the half site sequence at an affinity (118nM) similar to the full site PDREs (Figure 5A-B). Single substitutions of the half site sequence followed a similar trend to Pdr1p, with 5’-TTCCG<u>A</u>AA-3’ as the only single substitution to increase affinity (Sup. Figure 5C). The double substitution 5’-TTC<u>AA</u>GAA-3’ only reduced the Pdr3p affinity 2.5-fold, indicating that the binding of Pdr3p to the 18mer is less specific to the CGG triplet than Pdr1p (Sup. Figure 5D). Overall, these findings suggest that Pdr1p can bind to a single CGG triplet binding motif with high specificity, which explain the number of PDRE-free Pdr1p CUT&RUN peaks. In addition, these studies show that Pdr1p and Pdr3p half site binding prefers a downstream flanking sequence AA, indicating that the preference for flanking sequences for a single site is similar to that of a full PDRE site.

**Figure 5.**
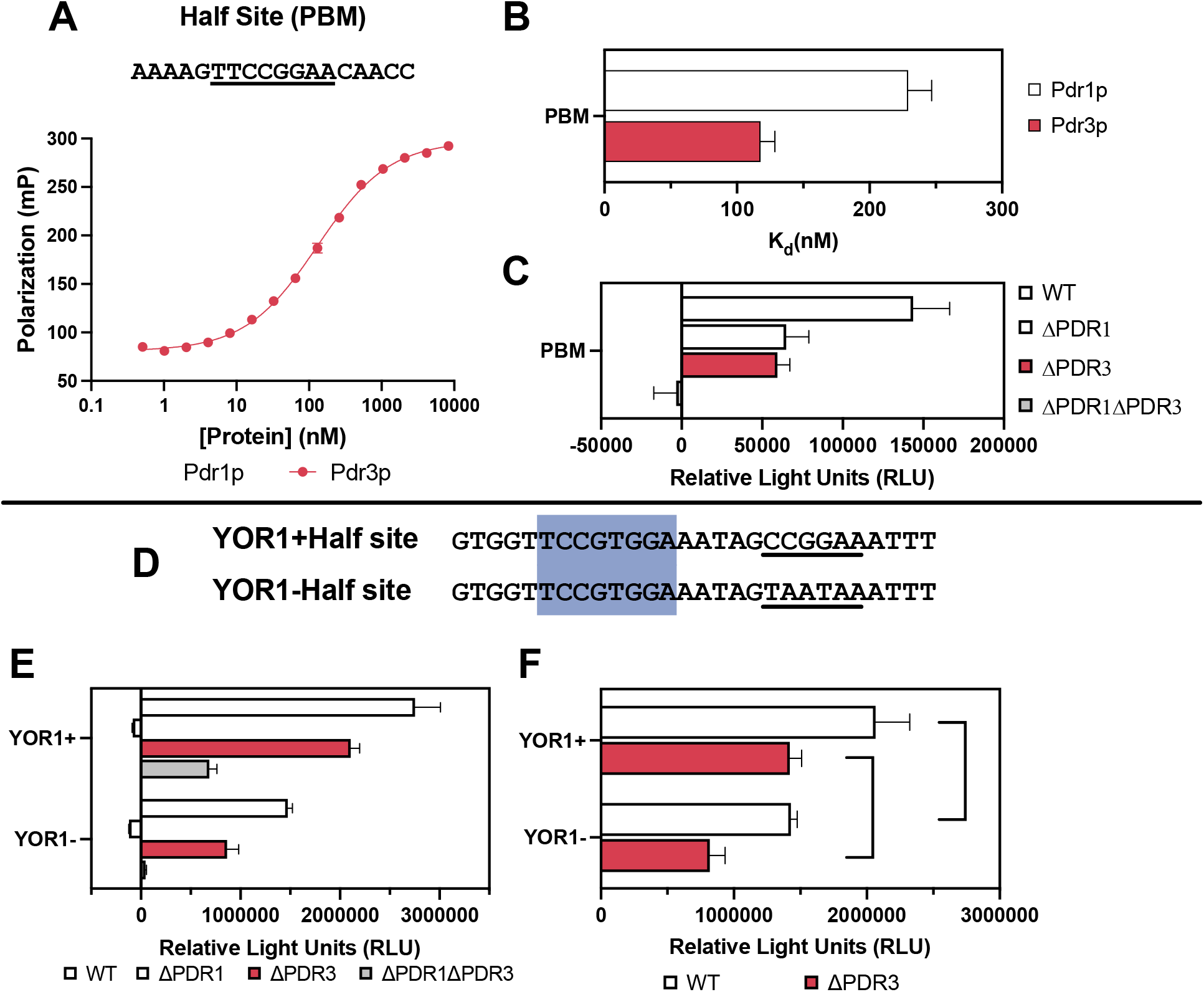
Pdr1p and Pdr3p DBDs bind the half site motif at similar affinities to the PDRE full sites, with half sites able to increase transcription from full sites. (**A**) Fluorescence polarization binding curves of a FAM-labeled 18mer PBM DNA containing 5’-TTCCGGAA-3’ and a 5mer flank incubated with a 2-fold serial dilution of either the Pdr1p or Pdr3p DBD. (**B**) Comparison of the dissociation constants of the half site motif to the Pdr1p and Pdr3p DBDs. Representative curves are shown for one experiment (three technical replicates). (**C**) Transcription induction of luciferase reporter plasmid containing the 18mer PBM DNA inserted into a minimal *CYC1* promoter. (**D**) The PDRE in the *YOR1* promoter (highlighted) has a neighboring half site motif (underlined). (**E**) Transcription induction of the WT *YOR1* PDRE and half site DNA, or with the half site removed. (**F**) Comparison of the impact of the *YOR1* half site sequence on transcription with nonspecific activity (ΔPDR1ΔPDR3) removed by subtracting its contribution to transcription. The observed differences in transcription between YOR1+ and YOR1- are statistically significant in both the WT and ΔPDR3 (Unpaired T test P values 0.013, and 0.002 respectively). Data points in panels B, C, E and F show standard deviations as error bars (n =3).

Since Pdr1p and Pdr3p demonstrated affinities for the half site PBM sequence similar to some full PDREs, the transcriptional activity of the PBM sequence was evaluated (Figure 5C). The results showed that the half site exhibited very weak transcriptional activity, reaching only 69% of the PDR5_4 site. This suggests that individual half sites are unlikely to significantly contribute to transcription in any promoter. To explore the possibility that half sites may enhance of the transcriptional activity of full PDRE sites, Pdr1 CUT&RUN peaks were scanned for PDREs in close proximity to the half site motif. The single PDRE in the *YOR1* promoter has a half site motif 5 nucleotides downstream (Figure 5D), and is a gene known to be regulated by Pdr1p and Pdr3p(24,30,58). The transcriptional activity of the DNA sequence corresponding to the *YOR1* PDRE and half site motif was tested, along with a sequence containing a mutated half site (Figure 5E). Since both DNA sequences had a large difference in nonspecific transcriptional activity (ΔPDR1ΔPDR3 strain), the Pdr1p and Pdr3p specific activity was compared (Figure 5F). There was significantly greater transcription in both the WT and ΔPDR3 strains for the *YOR1* sequence with an intact half site supporting the idea that half sites can contribute in some way to the activity of PDREs.

## DISCUSSION

For decades Pdr1p and Pdr3p have been recognized as the primary transcriptional regulators of many of the genes central to the *S. cerevisiae* PDR response. Over time it has been identified that despite their similarities, both Pdr1p and Pdr3p each have their own distinct activation mechanisms and regulons. Understanding of their DNA binding specificity has lagged behind, leading to identifying nearly identical binding motifs despite evidence of differing regulons. Here, we have clarified the impact of variation in the core PDRE and flanking sequences on both the DNA binding and transcriptional activity of Pdr1p and Pdr3p. Additionally, we have shown evidence for Pdr1p binding to DNA half sites both in vivo and in vitro.

Originally, we were surprised to see that only a small fraction of identified peaks for Pdr1p contained PDREs. Even when we used an irreproducible discovery rate (IDR) at a 0.05 cut-off to filter high confidence peaks among Pdr1p replicates, we only see 12.2% of peaks contain PDREs(59). While high signal peaks consistently contained multiple PDREs, the presence of a single PDRE was not strongly correlated with signal strength similar to what was previously reported in Fardeau et al.(48). Examination of the PDRE-free CUT&RUN peaks with MEME-ChIP identified half site motifs that were highly similar to the second ProBound binding mode and high affinity PDRE-free 8mers observed in previous PBM datasets. The Pdr1p PDRE-free peaks identified in CUT&RUN are likely not spurious but rather represent a different mode of binding. While attempting to refine the PDRE motif using promoters bound by Pdr1p from the YEASTRACT database, we noticed substantial inconsistency between published datasets (Sup. Figure 6)(48,60–65). We found the intersection between the Pdr1p CUT&RUN data, Fardeau et al. Pdr1p ChIP-seq, and Rossi et al. Pdr1p ChIP-exo represents the high confidence Pdr1p regulon (16 genes)(48,65). The CUT&RUN and Fardeau et al. datasets share an additional 4 genes (*RSB1*, *ERG27*, *VHR1*, *LAF1*) that also appear to have peaks in the raw data of the Rossi dataset but were not assigned(48,65). Overall, we see 114 shared genes between the Pdr1p CUT&RUN and ChIP-exo datasets, highlighting the increased sensitivity of these techniques over ChIP-seq (Sup. Figure 6). The high sensitivity of CUT&RUN allowed for identification of many additional genomic sites occupied by Pdr1p that corresponds to half site binding. We also compared our Pdr3p CUT&RUN data with the Rossi et al. Pdr3p ChIP-exo dataset. For both datasets Pdr3p was bound to substantially fewer promoters compared to Pdr1p(65). This limited occupancy by Pdr3p appears to be in agreement with prior work by Fardeau et al. that indicates Pdr1p and not Pdr3p was responsible for the early PDR response(48).

The data presented here reveals for the first time the importance of flanking sequences in PDRE function. The PDREs of the *PDR5* promoter provide a striking example of the impact of flanking sequences on PDRE dependent transcription with a 32-fold difference in activity between PDR5_2 and PDR5_4 (Figure 2D). By testing PDREs outside of their native promoter context, we were able to confirm that the previously reported weak activity of PDR5_4 is exclusively dependent on the binding site and a small flanking region(8,30). Pdr1p and Pdr3p DNA binding affinity are primarily dependent on sequence variation within the core 8bp PDRE sequence and the immediate 2bp flanking sequences, with further sequences having only a minor impact on affinity. Among the four PDRE types, both proteins favor different core sequence variation, with Pdr1p preferring A>B>D>C while Pdr3p prefers C>B>D>A (Figure 4). Interestingly, we observe that for PDRE dependent transcriptional activity that type B and C are driven equally by Pdr1p and Pdr3p while type A and D are primarily driven by Pdr1p and Pdr3p respectively. We propose that the variation in PDRE transcriptional activity is the result of both DNA affinity and DNA/protein complex conformation. Small variations in the conformation that Pdr1p and Pdr3p assume on DNA may alter the ability of their activation domains to recruit Mediator(66,67).

There are several other zinc cluster transcription factors that are part of the PDR regulatory network that also appear to bind an everted zero gap repeat similar to Pdr1p and Pdr3p, they but regulate a much narrower number of genes(9–11). Like Pdr1p and Pdr3p, limited biochemical data on these proteins has hampered the ability to identify their sequence specificities. One such protein is Rdr1p, which represses the transcription of five strongly induced PDR network genes(10). Rdr1p recognizes the type A and B PDREs in these promoters, yet it remains unclear why Rdr1p doesn’t regulate other PDRE containing promoters. We anticipate that in a similar fashion to Pdr1p and Pdr3p, these other PDR zinc cluster transcription factors likely have additional specificity for DNA binding based on flanking sequences that account for the smaller number of genes they regulate. Our in vitro binding experiments confirmed that the Pdr1p and Pdr3p DBDs can bind to a half site sequence discovered by both MEME and ProBound. The comparable affinity between the tested half site and full PDRE sequences indicates that more degeneracy is tolerated within the core binding motif than described in the four PDRE types. A half site alone was capable of only extremely weak transcription, indicating that presence of an individual half site is unlikely to confer regulation by either Pdr1p or Pdr3p in vivo (Figure 5). The presence of a half site in close proximity to a PDRE, (*YOR1* promoter), enhances transcriptional activity suggesting that half sites may function to increase occupancy of nearby PDREs by Pdr1p and Pdr3p. While individual half sites have low transcriptional activity, multiple half sites in a promoter may result in synergistic binding and transcriptional regulation, as seen in Rgt1p(68). Additionally, Pdr1p’s ability to bind half sites may have implications for its ability to heterodimerize with other zinc cluster transcription factors and recruit them to promoters with multiple half sites.

Based on the results presented here, we propose an updated model of Pdr1/3p transcriptional regulation. The core PDRE motif is necessary for regulation of a promoter by Pdr1/3p, while the flanking region and presence of nearby half sites heavily modulate the strength of regulation (Figure 6). This modulation helps to explain the variation in the strength of transcriptional regulation of different genes by Pdr1/3p despite the presence of the same type of PDREs. Additionally, differences in flanking regions alter the degree of regulation by Pdr1p and Pdr3p leading to transcription dependent on primarily one protein, or equal. We anticipate future work will elucidate the mechanism behind the function of these PDRE flanking regions.

**Figure 6.**
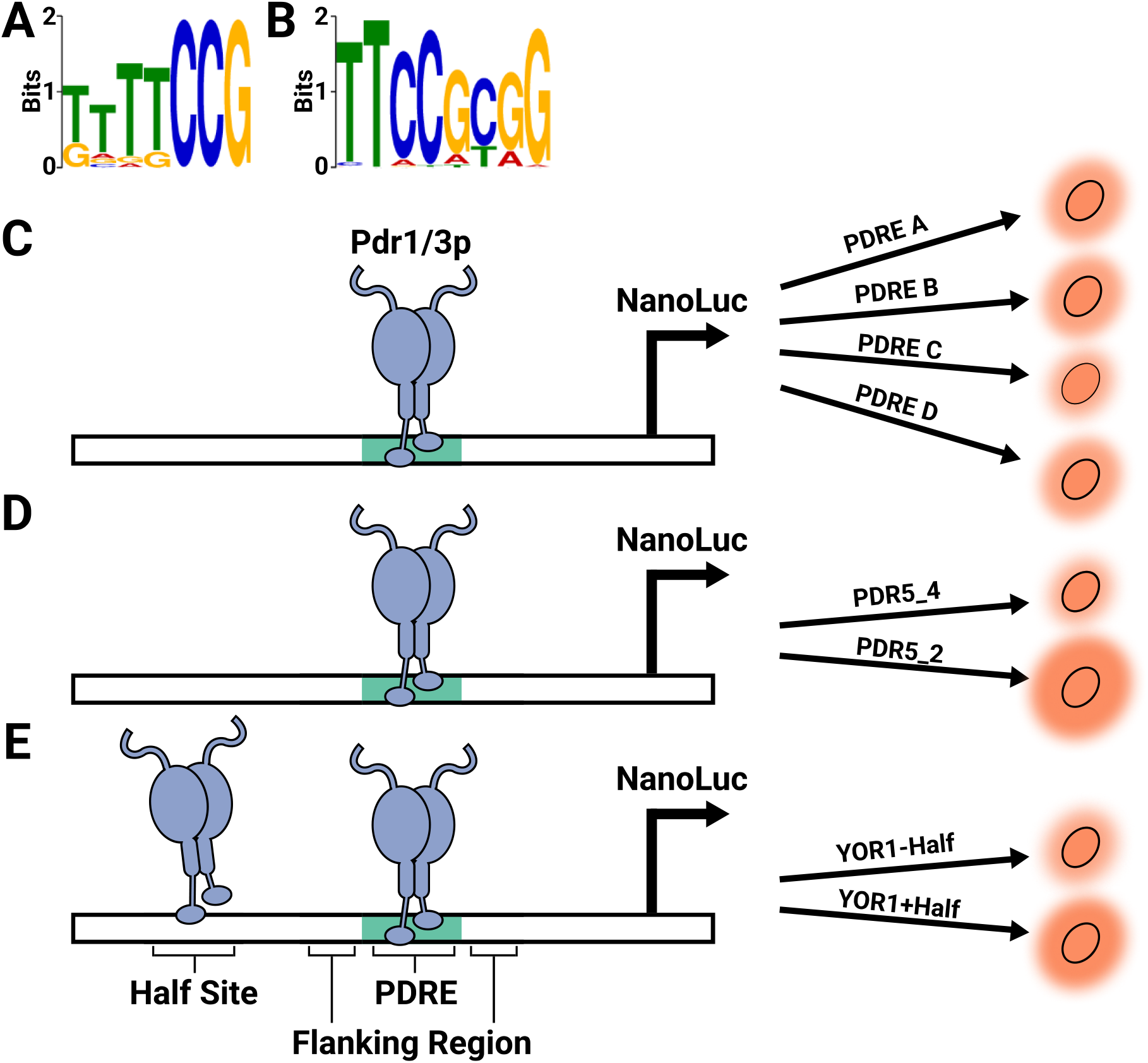
Summary of the impact of different DNA features on Pdr1p and Pdr3p dependent transcription. Motifs identified in the Pdr1p CUT&RUN data by STREME that correspond to the (**A**) half site and (**B**) PDRE binding. (**C**) Within the four PDRE types that capture variation within the core PDRE, only type C PDREs have weaker Pdr1/3p dependent transcription. (**D**) Sequence variation within the region flanking PDREs can dramatically alter Pdr1/3p dependent transcription, as seen with PDR5_2 and PDR5_4 PDREs. (**E**) The transcriptional activity of PDREs can be enhanced by nearby half sites, observed for the YOR1 PDRE.

## DATA AVAILABILITY

All raw and processed next generation sequencing data associated with this manuscript is available to download from the NCBI Gene Expression Omnibus repository(69) under the accession code GSE228788.

## SUPPLEMENTARY DATA

Supplementary Data are available at FEBS Letters online.

## ACKNOWLEDGEMENTS

We thank Karl Kuchler for the yeast strains and thank Claes Andréasson for plasmid pCA955. This work was supported by the Northwestern University NUSeq Core Facility. We also thank the Keck Biophysics and High-Throughput Analysis facilities for access to equipment.

## Abbreviations

PDR: pleiotropic drug resistance
PDRE: PDR responsive element
DBD: DNA binding domain
ABC: ATP binding cassette
PBM: protein binding microarray

