## Supplemental files for "Unraveling the Half and Full Site Sequence Specificity of the *Saccharomyces cerevisiae* Pdr1p and Pdr3p Transcription Factors"

**Supplemental Table 1.**

**CUT&RUN reveals Pdr1p binds to gene promoter regions that contain single or multiple type A, B, C, or D PDREs.** (**A**) Genes with a Pdr1p CUT&RUN peak between -1000 and +500 of the TSS, with genes sorted by the type of PDRE present in the peak. Genes in bold had Pdr1 peaks that contained multiple types of PDREs. Peak annotation was performed using the R package rGreat v1.99.5 in basalPlusExt mode.

| PDRE type | Genes with PDREs bound by Pdr1 | | | | |
| --- | --- | --- | --- | --- | --- |
| Type A 5'-TCCGCGGA-3' | ALD6 | CAP1 | CIS1 | CMC4 | DDP1 |
|  | GAC1 | GIP3 | HXT11 | HXT3 | HXT9 |
|  | ICY1 | IMA2 | **IMA3** | IMA4 | IML2 |
|  | IPT1 | ISU1 | LAC1 | **LAF1** | LDB7 |
|  | **MIG2** | NCE102 | **PDR15** | PDR16 | PDR3 |
|  | **PDR5** | PGA3 | RGI2 | RGL1 | RPS11A |
|  | RSB1 | SGA1 | SLA1 | SNF11 | **SNQ2** |
|  | SPO24 | VHR1 | XBP1 | YDR010C | YGP1 |
|  | YGR017W | YHI9 | YHR213W-A | YHR213W-B | YHR214W |
|  | **YMR103C** | YPL062W | YPL068C | YRR1 |  |
| Type B 5'-TCCACGGA-3' | ADY2 | ATG40 | BUD21 | CUE4 | DSF2 |
|  | ENA1 | ERG27 | GRE2 | **HXT8** | ICT1 |
|  | **IMA3** | **IMA5** | INA1 | LAM4 | LRP1 |
|  | MET28 | **MIG2** | MIM2 | MMO1 | MNN4 |
|  | MOD5 | MRP1 | OLE1 | PDR10 | **PDR5** |
|  | PHD1 | PXR1 | RLM1 | **RPL36B** | RPN4 |
|  | RPS0A | RSF2 | RSM10 | RTA1 | **RTS3** |
|  | SKI7 | SNC2 | **SNQ2** | SNR3 | SNR86 |
|  | SUR2 | SVF1 | TPO1 | TPO4 | TSL1 |
|  | TVS1 | YAP5 | YAP6 | YDR010C | YGR035C |
|  | YGR035W-A | YGR161W-C | YLR412C-A | YNCP0001C | YOR1 |
|  | YPL088W | YPL257W-A | YPL257W-B |  |  |
| Type C 5'-TCCGCGCA-3' | ARO9 | PUT4 | RPL27B | **RPL36B** | RPS18B |
|  | **SCS7** | **SNQ2** | SPL2 | TEF1 | UGO1 |
|  | YCK1 | YDR010C | YMR272W-B | YPR014C |  |
| Type D 5'-TCCGCGGG-3' | ARE2 | BSC2 | CAM1 | CDC34 | COA6 |
|  | CWP2 | DIG1 | DPI8 | FAA4 | HEM13 |
|  | **HXT8** | **IMA5** | **LAF1** | LSM7 | MRS3 |
|  | MRS6 | PDA1 | **PDR15** | PGA2 | PST1 |
|  | RTG3 | **RTS3** | **SCS7** | SFT2 | TYE7 |
|  | YDR274C | YGR161W-C | YKL096C-B | YKL097C | **YMR103C** |
|  | YMR272W-B |  |  |  |  |

**Supplemental Figure 1.**

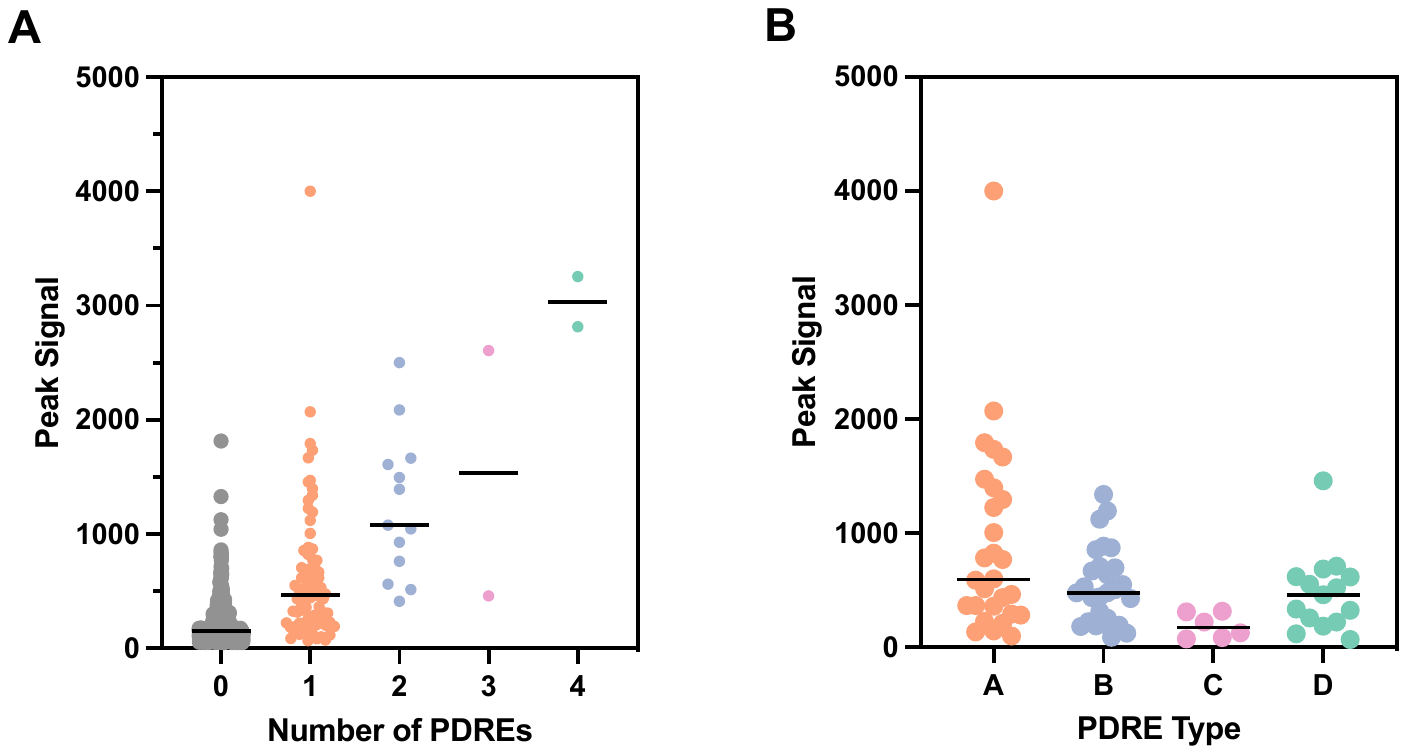

**Pdr1p genomic occupancy is dependent on the presence and type of PDREs.** (**A**) Pdr1p peaks identified by the peak caller MACS2 were sorted by number of type A-D PDREs. (**B**) Pdr1p peaks containing a single type A-D PDRE were sorted by type. The type C PDREs had significantly lower peak signal than type A, B, and D (Mann-Whitney U test *P* values 0.0018, 0.0069, and 0.0365, respectively. Peak signal corresponds to the signal score produced by MACS2.

**Supplemental Figure 2.**

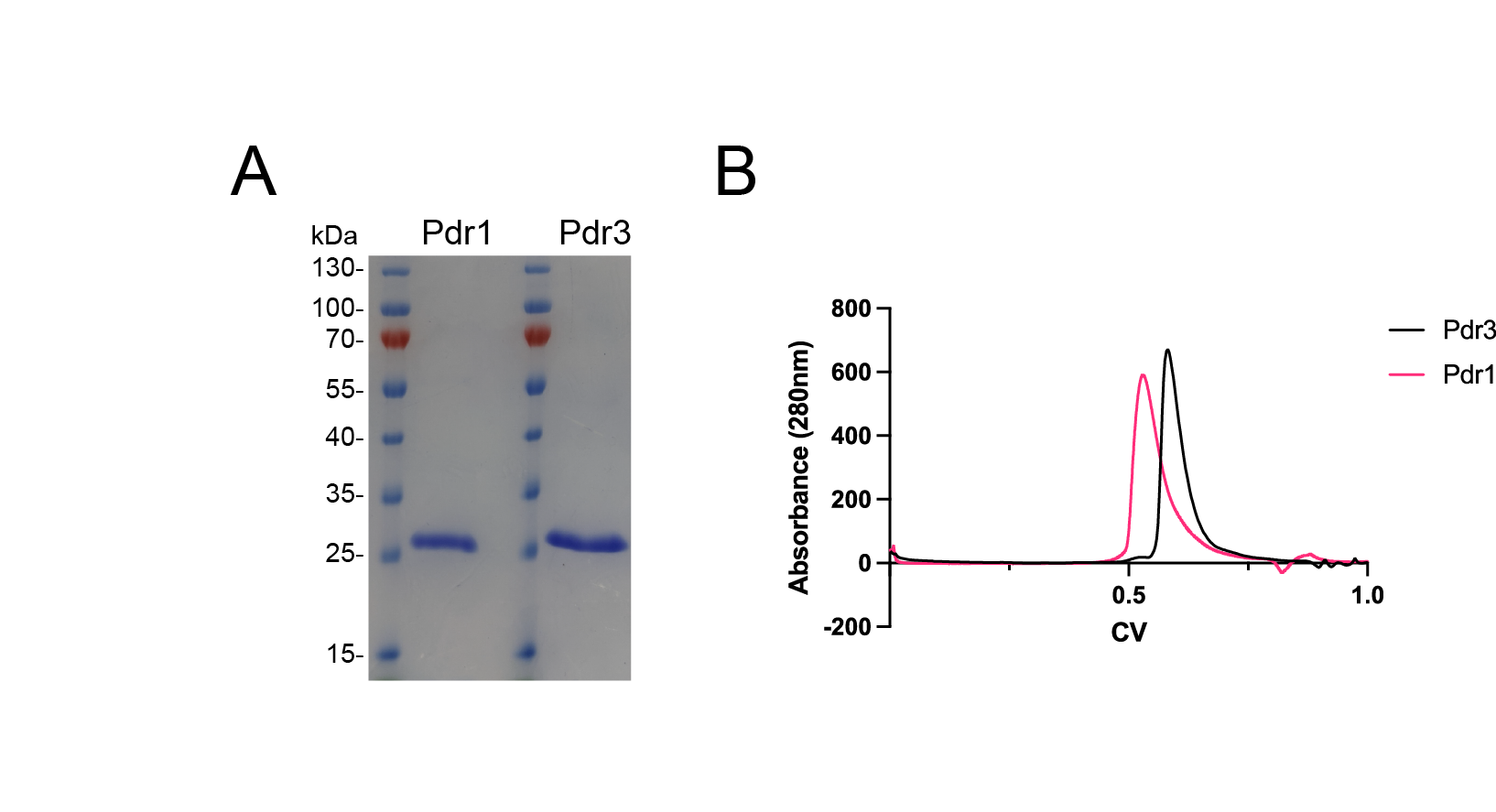

**Pdr1p and Pdr3p DBDs are pure and mono-disperse.** (**A**) SDS-PAGE gel of the Pdr1p and Pdr3p DBDs post purification indicates greater than 95% purity. (**B**) Size exclusion chromatography profiles of Pdr1p and Pdr3p DBDs injected onto a HiLoad 16/600 Superdex 200 pg column reflects a monodisperse dimer.

**Supplemental Figure 3.**

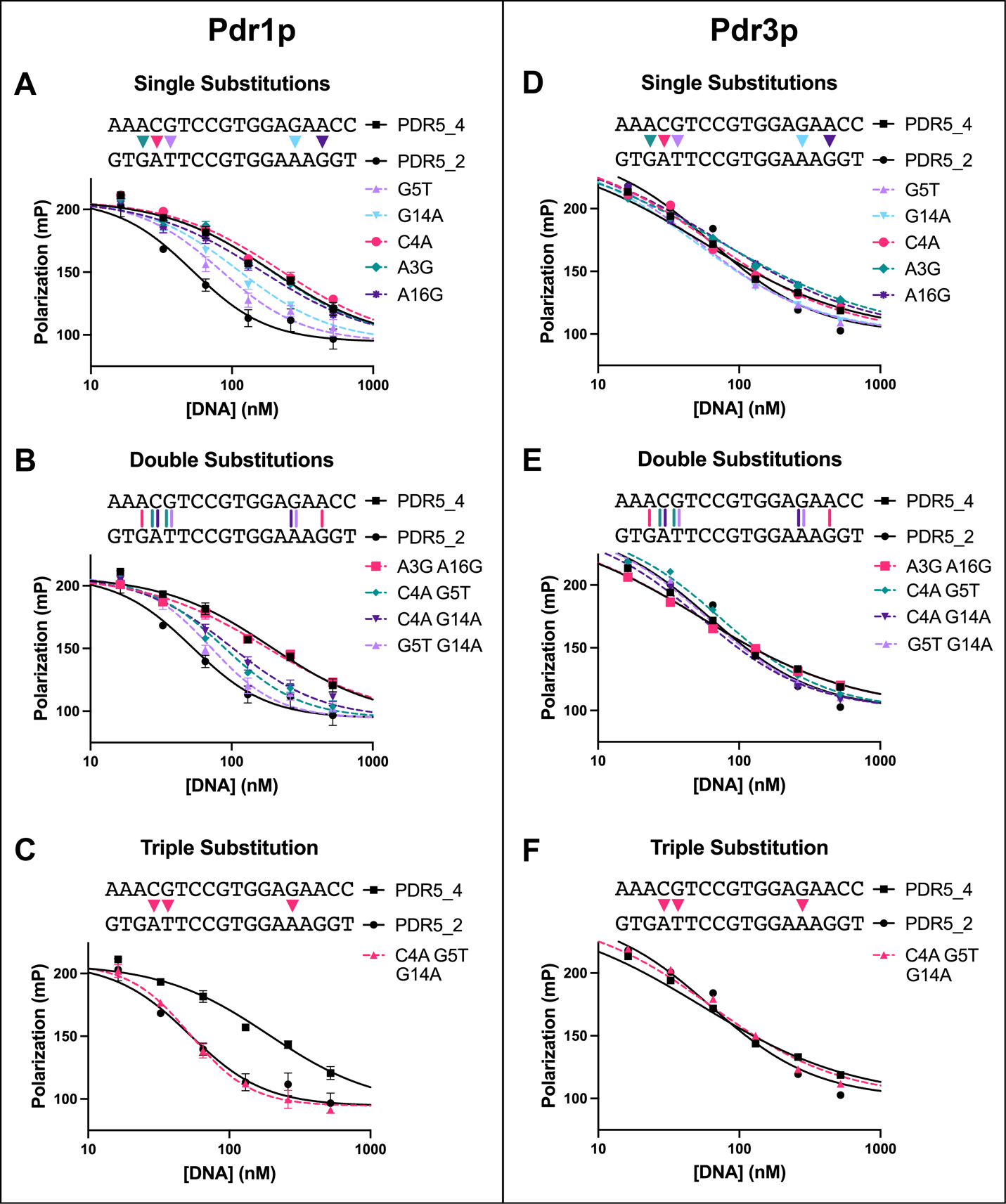

**The modification of the PDRE flanking sequences alone alters Pdr1p and Pdr3p DNA binding affinities.** The type B PDREs PDR5_2 and PDR5_4 were selected due to differences in both binding affinities and variable flanking sequences. Competitive fluorescence polarization binding curves of FAM-labeled PDR5_4 PDRE DNA and Pdr1p DBD incubated in the presence of unlabeled PDR5_4 PDRE DNA with either (**A**) single (**B**) double or (**C**) triple flanking base substitutions. Bases in the flanking sequences of the PDR5_4 PDRE were substituted with the corresponding base from PDR5_2. Competitive fluorescence polarization binding curves of FAM-labeled PDR5_4 PDRE DNA and Pdr3p DBD incubated in the presence of unlabeled PDR5_4 PDRE DNA with either (**D**) single (**E**) double or (**F**) triple flanking base substitutions. The data was analyzed in Graphpad Prism 9.0 using a sigmoidal 4PL fit with a shared top and bottom for all datasets. Representative curves are shown for one experiment (three technical replicates) and were repeated at least two times in triplicate.

**Supplemental Figure 4.**

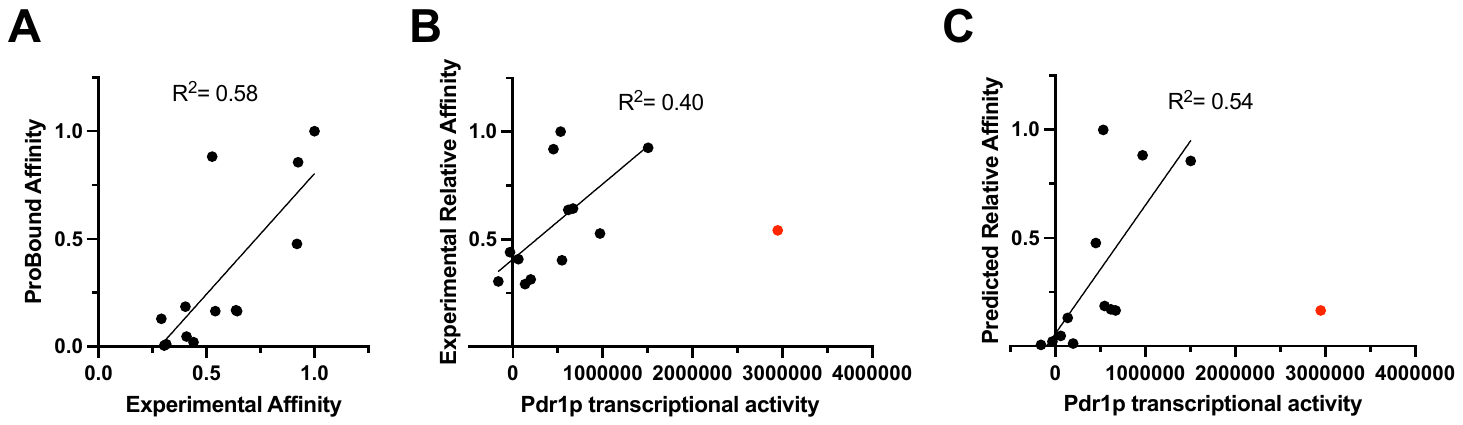

**Pdr1p DNA binding affinity has only a moderate impact on transcriptional activity.** (**A**) Relative affinities computed from experimental fluorescence polarization experiments, or from the ProBound PDRE binding mode. A relative affinity of 1 corresponds to the tightest bound DNA sequence. Comparison of the correlation between the (**B**) experimental binding affinities or (**C**) ProBound predicted affinities of Pdr1p to PDREs and the Pdr1p specific transcriptional activity of each PDRE. Pdr1p transcriptional activity was calculated by subtraction of the relative light units of the ΔPDR1ΔPDR3 strain from the ΔPDR3 strain for all PDREs. The data was analyzed in Graphpad Prism 9.0 using a linear regression, with the PDR5_2 PDRE (red) excluded.

**Supplemental Figure 5.**

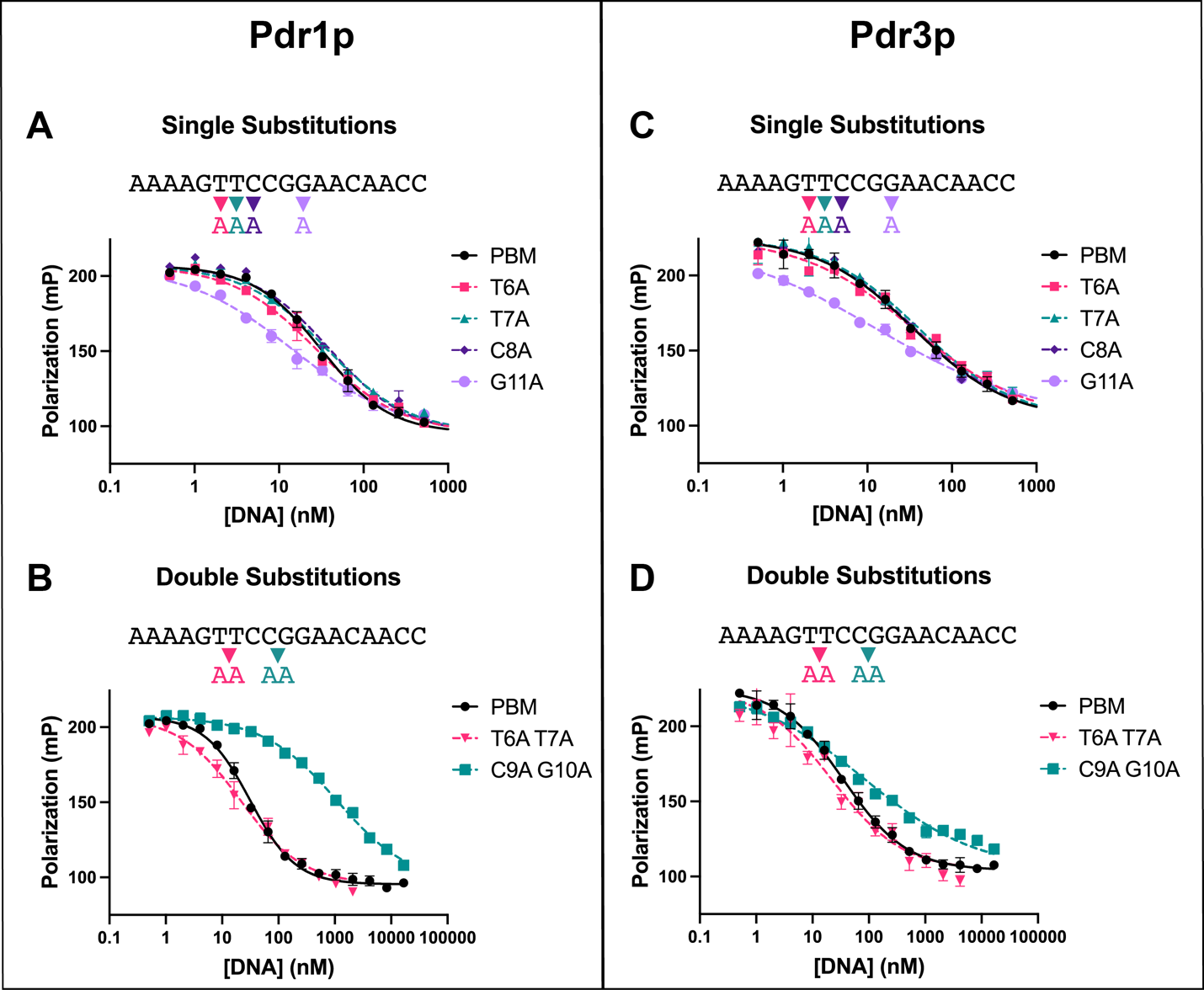

**Pdr1p and Pdr3p DBDs bind the half site motif is insensitive to changes in flanking nucleotides, with Pdr1p binding heavily dependent on the core CGG triplet**. Competitive binding curve of FAM-PBM with Pdr1p DBD incubated with unlabeled DNAs of the PBM sequence with either (**A**)single or (**B**) double base substitutions. Competitive binding curve of FAM-PBM with Pdr3p DBD incubated with unlabeled DNAs of the PBM sequence with either (**C**) single or (**D**) double base substitutions. The data was analyzed in Graphpad Prism 9.0 using a sigmoidal 4PL fit with a shared top and bottom for all datasets. Representative curves are shown for one experiment (three technical replicates) and were repeated at least two times in triplicate.

**Supplemental Figure 6.**

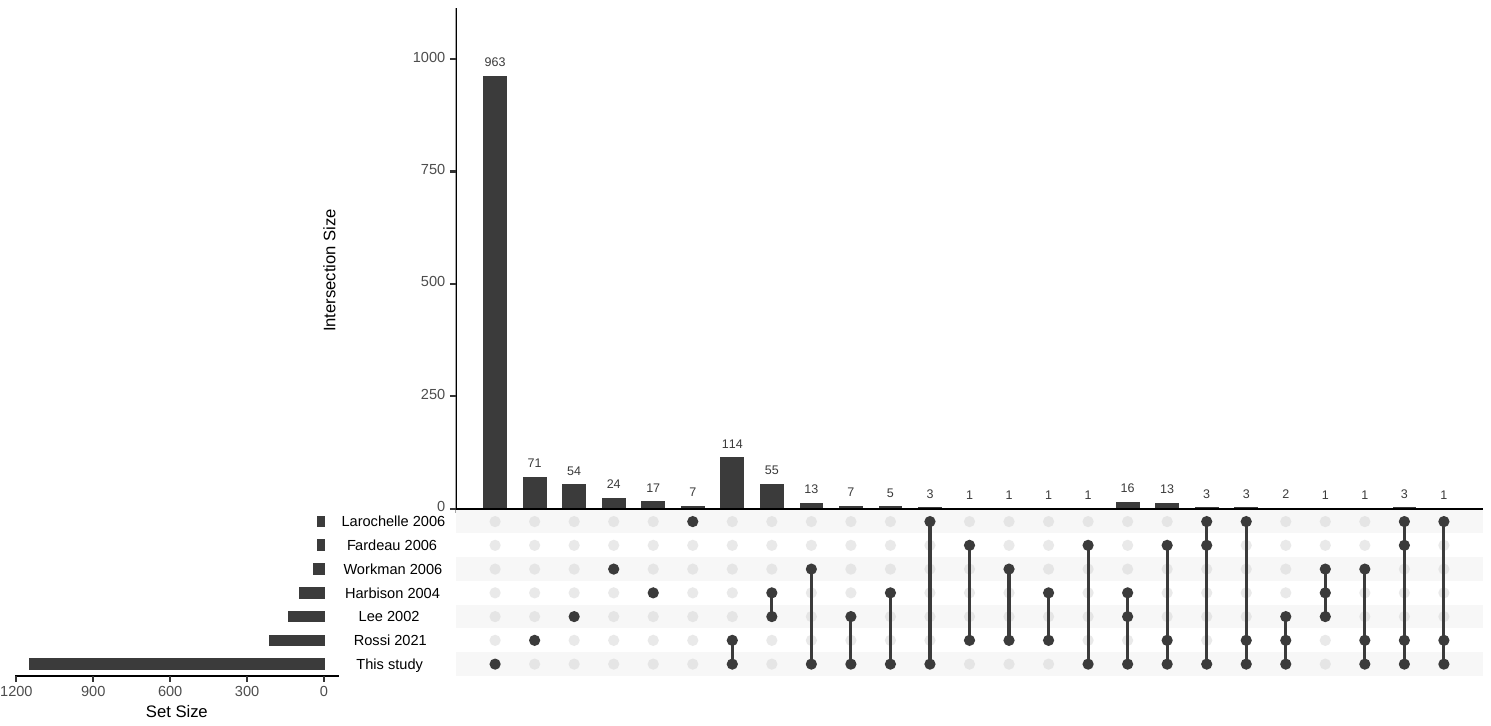

**CUT&RUN identifies more genes bound by Pdr1p than previous ChIP experiments.** (**A**) UpSet plot of the intersection of genes bound by Pdr1p in different published ChIP-chip, ChIP-seq, and ChIP-exo datasets compared to the current CUT&RUN dataset. Gene lists were obtained from either the original publication or YEASTRACT and UpSet plot was produced using the R package UpSetR v1.4.0.
